# A Computational Algorithm for Optimal Design of Bioartificial Organ Scaffold Architectures

**DOI:** 10.1101/2024.04.16.589695

**Authors:** Martina Bukač, Sunčica Čanić, Boris Muha, Yifan Wang

## Abstract

We develop a computational algorithm based on a diffuse interface approach to study the design of bioartificial organ scaffold architectures. These scaffolds, composed of poroelastic hydrogels housing transplanted cells, are linked to the patient’s blood circulation via an anastomosis graft. Before entering the scaffold, the blood flow passes through a filter, and the resulting filtered blood plasma transports oxygen and nutrients to sustain the viability of transplanted cells over the long term. A key issue in maintaining cell viability is the design of ultrafiltrate channels within the hydrogel scaffold to facilitate advection-enhanced oxygen supply ensuring oxygen levels remain above a critical threshold to prevent hypoxia. In this manuscript, we develop a computational algorithm to analyze the plasma flow and oxygen concentration within hydrogels featuring various channel geometries. Our objective is to identify the optimal hydrogel channel architecture that sustains oxygen concentration throughout the scaffold above the critical hypoxic threshold.

The computational algorithm we introduce here employs a diffuse interface approach to solve a multi-physics problem. The corresponding model couples the time-dependent Stokes equations, governing blood plasma flow through the channel network, with the time-dependent Biot equations, characterizing Darcy velocity, pressure, and displacement within the poroelastic hydrogel containing the transplanted cells. Subsequently, the calculated plasma velocity is utilized to determine oxygen concentration within the scaffold using a diffuse interface advection-reaction-diffusion model. Our investigation yields a scaffold architecture featuring a hexagonal channel network geometry that meets the desired oxygen concentration criteria. Unlike classical sharp interface approaches, the diffuse interface approach we employ is particularly adept at addressing problems with intricate interface geometries, such as those encountered in bioartificial organ scaffold design. This study is significant because recent developments in hydrogel fabrication make it now possible to control hydrogel rheology [20, 14], and utilize computational results to generate optimized scaffold architectures.

**MSC codes:** 76S05; 76-04; 76D05; 92-10; 92-04

## 1. Introduction

Bioartificial organ is an engineered device made of living cells and a biocompatible scaffold that can be implanted or integrated into a human body– interfacing with living tissue–to replace a natural organ, or to duplicate or augment a specific organ function [29]. Biocompatible scaffold is a biocompatible base material in which cells and growth factors are embedded to construct a substitute tissue, which can be used in e.g. bioartificial organ design [20]. For example, a biocompatible scaffold used in the bioartificial pancreas design is a biocompatible hydrogel (e.g. agarose gel) which houses transplanted pancreatic islets (conglomerations of pancreatic cells) that contain the insulin-producing *β*-cells. Hydrogels are hydrophilic polymers that are poroelastic, and that have the ability to absorb a large volume of fluid, which makes them particularly suitable materials for biomedical applications. In bioartificial pancreas design presented in [26], the biocompatible hydrogel containing the insulin-producing cells is encapsulated in a nano-pore, semi-permeable membrane (filter), and the resulting bioartificial organ is connected to the host’s cardiovascular system via an anastomosis graft (a tube) for the advection-enhanced nutrients delivery to the cells, and insulin distribution away from the cells. See Figure 1.

**Fig. 1.**
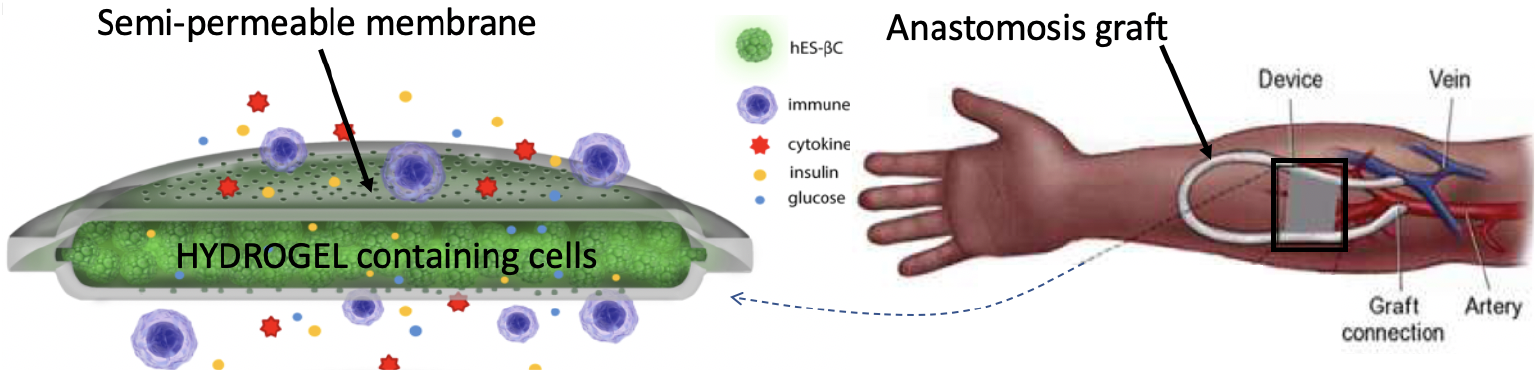
Left: Schematic illustration of immunoprotective encapsulation device containing human embryonic stem cell-derived *β* clusters (hES-*β*C). Right: Schematic illustration showing anastomosis graft connecting encapsulation device to the host vasculature. Courtesy of Dr. S. Roy (Bioengineering, UCSF).

A major challenge in the encapsulation-based bioartificial organ design is maintaining the survival of transplanted cells within the bioartifical organ scaffold for an extended period of time. In particular, long-term viability of pancreatic cells is directly affected by the sufficient access to nutrients for survival, of which oxygen is the limiting factor [12].

One of the goals of this manuscript is to develop the mathematical and computational approaches to investigate oxygen supply to the transplanted cells by studying hydrogel architecture and optimal design of ultrafiltrate channels within the encapsulated poroelastic hydrogels for advection-enhanced reaction-diffusion processes of oxygen transport to the cells. This is significant because recent developments in hydrogel fabrication make it now possible to control hydrogel elasticity and hydrogel rheology using, e.g., the approaches employed and reviewed in [20, 14].

In this manuscript we study oxygen-carrying blood plasma flow and oxygen concentration in three different scaffold architectures, whose geometries are inspired by different biological flow-nutrients scenarios. The first geometry consists of vertical ultrafiltrate channels drilled through a hydrogel, which is the simplest, and a standard procedure used in the design of bioartificial pancreas [19, 14, 27]. The second geometry consists of the branching channels, inspired by the architecture of the branching vessels in the human body. The third geometry consists of the hexagonal channels. This was inspired by the biological (epithelial) tissues in which interstitial fluid flows through a network of irregularly arranged interstices between hexagonally shaped cells, which supports their structural and functional integrity [9]. To work with comparable fluid flow scenarios, the three geometries are generated so that the total fluid channels’ volume (i.e., the area of the 2D channels) is the same in all three geometries.

We are interested in a scaffold architecture with the geometry that provides concentration of oxygen that is as uniform throughout the scaffold as possible and above a minimum concentration below which hypoxia occurs. For this purpose we introduce two sets of models, one for the fluid flow and one for oxygen concentration in the scaffold. The scaffold is modeled as a 2*D* domain containing a network of channels separating the regions of poroelastic medium describing a hydrogel. See Figure 2 below. In the last part of the manuscript we introduce a 3*D* computational model to show that for the “optimal” geometry, the 2*D* simulations capture the main features of flow and oxygen concetration. The flow of blood plasma through the network of channels is modeled by the time dependent Stokes equations for an incompressible, viscous fluid. The channel flow is coupled to the filtrating flow in the poroelastic hydrogel, which is modeled by the Biot equations [31]. Two sets of coupling conditions at the interface between the channel flow and hydrogel poroelastic medium are employed: the kinematic and dynamic coupling conditions. The kinematic coupling conditions describe continuity of normal (vertical to the interface) velocity and a slip in the tangential component of the velocity, known as the Beaver-Joseph-Saffman condition. The dynamic coupling conditions state the balance of forces at the interface. Linear coupling is considered, i.e., the coupling conditions are evaluated at a reference interface. Subsequently, the resulting fluid velocity is used as an input data in an advection-reaction-diffusion model for oxygen concentration within the scaffold.

**Fig. 2.**
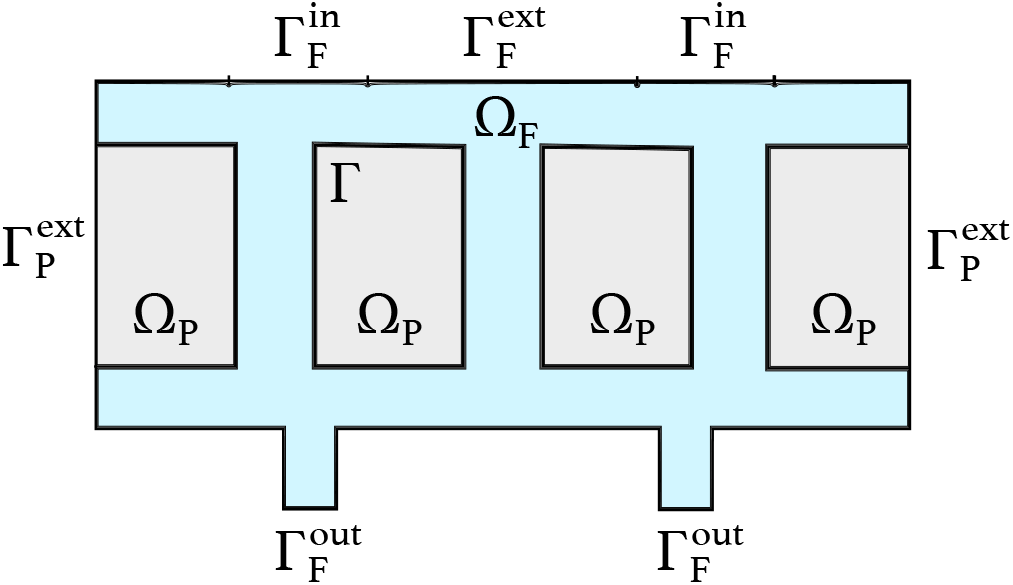
*An example of a domain* Ω = Ω_*F*_ ∪ Ω_*P*_ *demonstrating the two inlets* 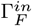 *at the top and the two outlets* 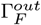 *at the bottom*.

To solve the coupled problems computationally, we implemented a diffuse interface approach [1, 13, 28]. The diffuse interface approach has several advantages over the classical sharp interface approach. In the diffuse interface approach, the mesh nodes do not have to be aligned with the interface, whose location is now captured using a phase-field function. Instead, the problem unknowns are defined on the entire domain, and the mesh remains fixed. This is particularly useful in cases where the interface between the two regions is difficult to determine exactly (e.g. “fuzzy” boundaries between regions obtained from medical images), or when the geometry of the interface is complex, leading to inaccurate approximations of the boundary integrals. Furthermore, advantages of this approach are apparent in problems with moving geometries and in the development of geometric optimization algorithms for optimal channel architecture design, which we plan to do next. In these cases a diffuse interface approach avoids the creation of complicated meshes following optimizing scaffold geometries, which would be the case in the classical sharp interface approaches. In this work we develop a diffuse interface approach to solve both the fluid and oxygen problems. In both cases, a finite element method is used for spatial discretization, and a backward Euler method for time stepping. We perform a validation of our numerical diffuse interface approach by comparing the solutions of the diffuse interface problem with the already validated solutions of a sharp interface approach, showing excellent agreement. Additionally, to justify the computationally less expensive 2*D* simulations, for the optimal geometry we perform the full 3*D* simulations and show that the 2*D* simulations we used to find the optimal geometry, capture all the important features of plasma flow and oxygen concentration in the corresponding 3*D* scaffold.

The diffuse interface approach is then used to investigate flow, pressure, and oxygen concentration in the three main geometries, discussed above. As mentioned earlier, the diffuse interface approach developed here is particularly suitable from the computational standpoint to study a number of different interface geometries. We find that the ultrafiltrate channel network with the hexagonal channel geometry provides the best solution in terms of oxygen concentration distribution and magnitude. Oxygen concentration in the hexagonal case is most uniformly distributed throughout the entire scaffold, and the values of concentration are all well above the critical value of oxygen below which hypoxia occurs. This is not the case for the other two geometries considered here (vertical channels and branching channels). We then further investigate the reasons why the hexagonal geometry performs best, and conclude that the relative angle of the channels with respect to the flow direction is one of the main contributing factors to higher Darcy velocity within the poroelastic hydrogel, and consequently the higher concentration values in between the ultrafiltrate channels. The results obtained in this manuscript can be used in optimal scaffold design by implementing hydrogel manufacturing techniques recently developed in [20, 14].

This manuscript is organized as follows. In Section 2 we introduce the model equations in strong/differential and weak/integral formulations. In Section 3 the diffuse interface method is discussed, and the fluid and oxygen problems are presented in the weak (integral) diffuse interface formulations. Numerical discretization based on a finite element approach is also discussed in this section. The computational setting, which includes the computational geometries described by the phase-field function, and parameter identification, is presented in Section 4. In this section we also show the validation of the numerical solver by comparing the diffuse interface solution with a monolithic solver utilizing a sharp-interface approach. Numerical results are presented in Section 5. In Section 6, we present 3*D* simulations of the optimal geometry demonstrating strong agreement with the 2*D* simulations. Conclusions are presented in Section 7.

## 2. Mathematical model

To model the entire scaffold, we introduce a bounded domain Ω ⊂ ℝ^*d*^, with *d* = 2, 3, consisting of a fluid region, Ω_*F*_, corresponding to the blood or blood plasma flow region bringing oxygen and nutrients to the transplanted cells, and a poroelastic solid region, Ω_*P*_, corresponding to the poroelastic medium such as a biocompatible hydrogel, which hosts the transplanted cells. The entire bioartificial organ scaffold is then comprised of the two regions so that 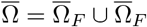 and Ω_*F*_ ∩ Ω_*P*_ = ∅ (see Figure 2). The fluid flowing through the channels comprising the region Ω_*F*_ enters the poroelastic medium Ω_*P*_ across the interface Γ, which is the interface between Ω_*F*_ and Ω_*P*_, defined by 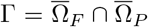. Figure 2 shows a sketch in which the fluid domain Ω_*F*_ consists of two horizontal channels corresponding to the “inlet” and “outlet” channels, namely, they contain the inlet boundary 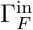 and the outlet 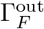, respectively. The two horizontal channels are connected via three vertical channels describing the simplest scaffold architecture which has been used in several research studies on bioartificial organ design [19, 14, 27]. The vertical ultrafiltrate channels are drilled to enhance advection-dominated oxygen and nutrients supply to the cells residing in Ω_*P*_ . This is just one, and the simplest example of the scaffold architecture that we will consider in this manuscript.

To model the interaction between the flow of blood plasma in Ω_*F*_ and the filtration of blood plasma through Ω_*P*_, we will use a fluid-poroelastic structure interaction model, which we present next.

### 2.1 Fluid-poroelastic structure interaction

We assume that Ω_*F*_ contains a viscous, incompressible, Newtonian fluid, such as blood plasma, whose dynamics can be described by the Stokes equations:

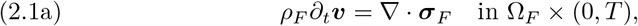

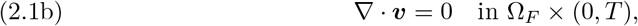

where ***v*** stands for the fluid velocity, *ρ*_*F*_ is density, and ***σ***_*F*_ is the Cauchy stress tensor defined by:

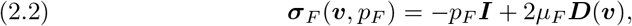

where *p*_*F*_ represents the fluid pressure, *µ*_*F*_ is the fluid viscosity, and ***D***(∇***v***) = (∇***v*** + ∇***v***^*τ*^)*/*2 is the strain rate tensor. Namely, equation (2.2) is a constitutive law describing a Newtonian fluid such as blood plasma.

The poroelastic medium, such as agarose gel used in the design of a bioartificial pancreas, see [12], can be described by the well-known Biot model, which has been used to describe hydrogels in [30]. The Biot model consists of the following equations:

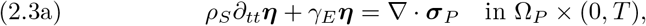

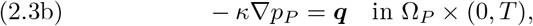

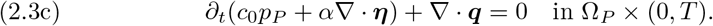

Equation (2.3a) describes the elastodynamics of the elastic skeleton, i.e., the solid phase in the Biot model, and is given in terms of the displacement of the elastic skeleton, denoted by ***η***, from its reference configuration Ω_*P*_ . The total Cauchy stress tensor ***σ***_*P*_ for the poroelastic region is defined by

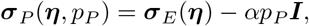

where *p*_*P*_ is the fluid pore pressure, *α* is the Biot-Willis parameter accounting for the coupling strength between the fluid and solid phases, and ***σ***_*E*_ denotes the elastic stress tensor, described by the Saint Venant-Kirchhoff constitutive model as:

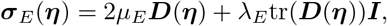

where *λ*_*E*_ and *µ*_*E*_ are Lamé parameters. In the elastodynamics equation (2.3a) above, constant *ρ*_*S*_ denotes density of the poroelastic matrix, and *γ*_*E*_ is a spring coefficient which accounts for the three-dimensional structure of the material holding different components together. Since our numerical simulations are performed in 2*D*, adding the term *γ*_*E*_***η*** accounts for the three-dimensional elastic energy effects and helps to avoid spurious numerical solutions in which sections of Ω_*P*_ might be pulled in the direction of vertical fluid flow allowing unreasonably large vertical displacements that do not occur in realistic 3*D* models due to the presence of an elastic restoring force in the third spatial dimension. Equation (2.3b) in the Biot model is the Darcy’s law and equation (2.3c) represents mass conservation.

#### Coupling conditions

To couple the fluid flow model with the Biot poroelastic medium model we use a set of four coupling conditions: two kinematic coupling conditions and two dynamic coupling conditions, which are evaluated at the location Γ of the interface between the two models. To state the coupling conditions introduce ***n***_*F*_ to denote the outward unit normal vector to *∂*Ω_*F*_ . At the interface Γ the following coupling conditions hold:

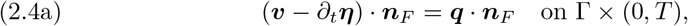

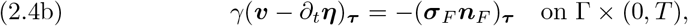

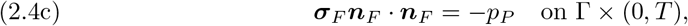

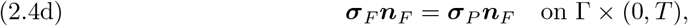

where for a vector ***v***, we define a projection onto the local tangent plane on Γ as:

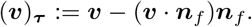

The first two conditions provide information about kinematic quantities such as velocities (kinematic coupling conditions), and the second two conditions provide information about coupling of stresses/forces (dynamic coupling conditions). More precisely, equation (2.4a) describes the fluid mass conservation across the interface, i.e., the normal components of the free fluid velocity ***v*** relative to the velocity of motion of the poroelastic matrix *∂*_*t*_***η*** is equal to the filtration velocity ***q*** across the interface. Equation (2.4b) describes the Beavers-Joseph-Saffman condition (2.4b) with slip rate *γ >* 0, namely, the tangential component of the free fluid velocity slips at the interface with the slip rate proportional to the fluid shear stress (***σ***_*F*_ ***n***_*F*_)_***τ***_ . Equations (2.4c) and (2.4d) describe the continuity of pressures and total stresses at the interface.

#### Boundary and initial conditions

Problem (2.1a)-(2.4d) is supplemented with boundary and initial conditions. In our notation, we represent the exterior boundary as Γ^*ext*^ = *∂*Ω. The exterior boundary is divided into three distinct parts: the fluid inflow boundary, the fluid outflow boundary, and the impenetrable part of the boundary, 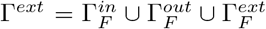. For the fluid problem, we impose the following boundary conditions:

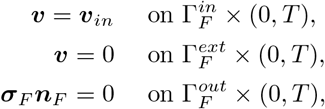

where ***v***_*in*_ is a prescribed velocity, specified in Section 4.

For the poroelastic structure, we impose no flow and zero displacement at the external boundaries:

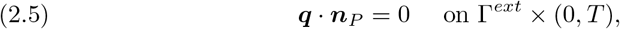

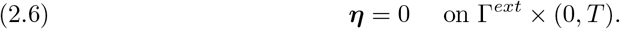

Initially, we assume that the fluid is at rest and that the deformable poroelastic structure is in its reference configuration. Thus, we have:

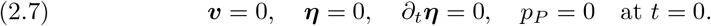

#### Weak formulation

To define the weak formulation of problem (2.1a)-(2.7) we introduce the following function spaces. Given an open set *S*, we consider the usual Sobolev spaces *H*^*k*^(*S*), with *k* ≥ 0, and introduce the following function spaces:

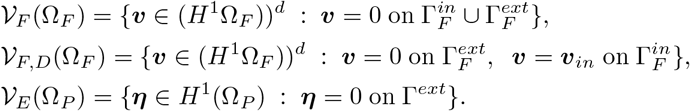

Here *𝒱*_*F*_ (Ω_*F*_) corresponds to the test space for the fluid velocity, *𝒱*_*F,D*_(Ω_*F*_) is associated with the solution space for the fluid velocity, and *𝒱*_*E*_(Ω_*P*_) is associated with the function space for the poroelastic matrix discplacement. The spaces *H*^1^(Ω_*P*_) and *L*^2^(Ω_*F*_) are associated with the solutions spaces for the fluid pressures in Ω_*p*_ and Ω_*F*_ respectively.

We say that (***v***, *p*_*P*_, ***η***, *p*_*F*_) ∈ *L*^2^(0, *T* ; *𝒱*_*F,D*_(Ω_*F*_) × *H*^1^(Ω_*P*_) × *L*^∞^(0, *T* ; *𝒱*_*E*_(Ω_*P*_)) × *H*^−1^(0, *T* ; *L*^2^(Ω_*F*_) is a weak solution if for every (***w***, *ψ*_*P*_, ***ζ***, *ψ*_*F*_) ∈ *𝒱*_*F*_ (Ω_*F*_) ×*H*^1^(Ω_*P*_) ×*𝒱*_*E*_(Ω_*P*_) × *L*^2^(Ω_*F*_), the following equality is satisfied in *𝒟* ^′^ (0, *T*):

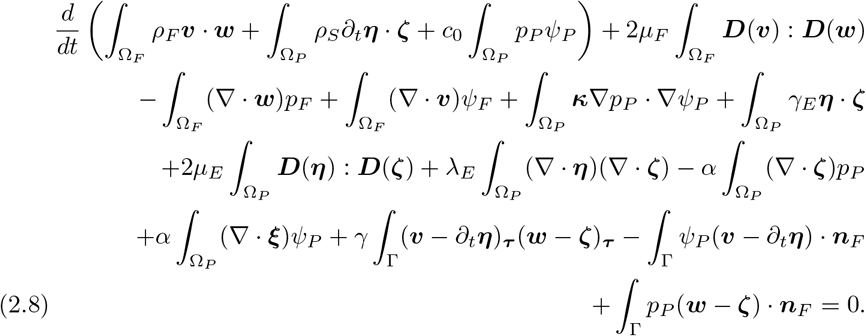

We note that the weak formulation is obtained using the primal formulation for the Biot problem, i.e., equations (2.3b) and (2.3c) have been combined so that equation (2.3c) is written only in terms of *p*_*P*_ and ***η***. The Darcy velocity can be computed by postprocessing using (2.3b).

Once the fluid velocity ***v*** and Darcy velocity ***q*** are computed from the fluidporoelastic structure interaction problem specified above, we can use this velocity information to formulate an advection-reaction-diffusion problem describing oxygen transport in the bioartificial organ consisting of ultrafiltrate channels Ω_*F*_ and the poroelastic medium Ω_*B*_ containing the cells.

### 2.2 Advection-reaction-diffusion

To model the transport of oxygen, we use an advection-reaction-diffusion model describing oxygen transport in the fluid domain Ω_*F*_ and in the poroelastic medium Ω_*P*_ . Oxygen transport in the human vascular system and tissues has been studied by many authors [4, 5, 17, 23, 22], and we adopt the approach from [5] to study oxygen transport in the scaffold:

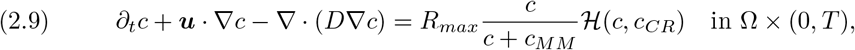

where *c* is concentration of oxygen, and *D* is a diffusion coefficient equal to *D*_*F*_ in Ω_*F*_ and to *D*_*P*_ in Ω_*P*_ . The velocity ***u*** is equal to ***v*** in Ω_*F*_, and to ***q*** in Ω_*P*_ . The reaction term on the right-hand side is active in Ω_*P*_ only and it accounts for the consumption of oxygen in the scaffold. In particular, this reaction term depends on the maximum oxygen consumption rate, *R*_*max*_ *<* 0, the Michaelis-Menten constant, *c*_*MM*_, corresponding to the oxygen concentration when the consumption rate drops to 50% of its maximum [5], and the function *ℋ* (*c, c*_*CR*_), defined by

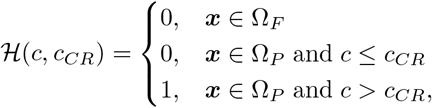

accounts for the regions of the tissue/poroelastic matrix where the oxygen concentration falls below a critical concentration *c*_*CR*_ below which necrosis is assumed to occur [5, 6]. The values of the parameters are all specified in [5, 6].

#### Boundary and initial conditions

Equation (2.9) is supplemented with the following boundary conditions:

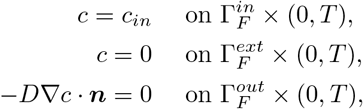

where ***c***_*in*_ is a given quantity, specified in Section 4. Thus, these boundary conditions say that we have a prescribed oxygen concentration at the inlet, zero oxygen concentration at the top boundary of the scaffold between the inlet regions, and zero diffusive oxygen flux at the outlet.

Initially, the concentration is set to zero:

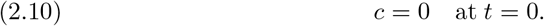

#### Weak formulation

To write the weak formulation of problem (2.9)-(2.10) we introduce the following function spaces:

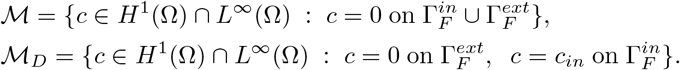

Here *ℳ* is associated with the test space for oxygen concentration and *ℳ*_*D*_ is associated with the solution space for *c*.

We say that *c* ∈ *L*^2^(0, *T* ; *ℳ*_*D*_) is a weak solution if *c* ≥ 0 and if for every *s* ∈ *ℳ*, the following equality is satisfied in *𝒟* ^′^ (0, *T*):

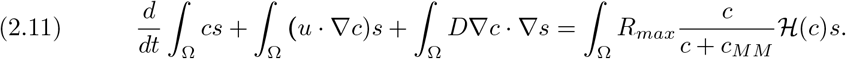

We have now specified two problems: a fluid-poroelastic structure interaction problem and an advection-reaction-diffusion problem, that we would like to solve for the fluid velocity and poroelastic structure displacement, and for oxygen concentration. The plan for this manuscript is to investigate three different scaffold architectures, motivated by biological structures, and numerically test which one provides the scaffold architecture with oxygen concentration that is closest to the uniform distribution of oxygen and is above the known minimal value *c*_*opt*_ for which uninhibited insulin production by the *β*-cells is guaranteed [25]. See Section 5. For this purpose we developed two numerical methods, one for the fluid-poroelastic structure interaction problem, and one for the advection-reaction-diffusion problem, which we describe next.

## 3. Numerical method

To solve the fluid-poroelastic structure interaction and transport problems numerically, we use the diffuse interface method [1, 13]. Let *χ* denote the Heaviside function which equals one in Ω_*F*_ and zero in Ω_*P*_ . Let a phasefield function Φ : Ω → [0, 1] be a regularization of the Heaviside function such that Φ ≈ 1 in Ω_*F*_, Φ ≈ 0 in Ω_*P*_, and Φ smoothly transitions between these two values on a “diffuse” layer of width *E* (see Figure 3). We suppose that *d*Γ ≈ |∇Φ|*dx* and 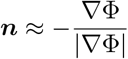. Using this notation, for functions *F* and *f* defined on Ω, we can write:

**Fig. 3.**
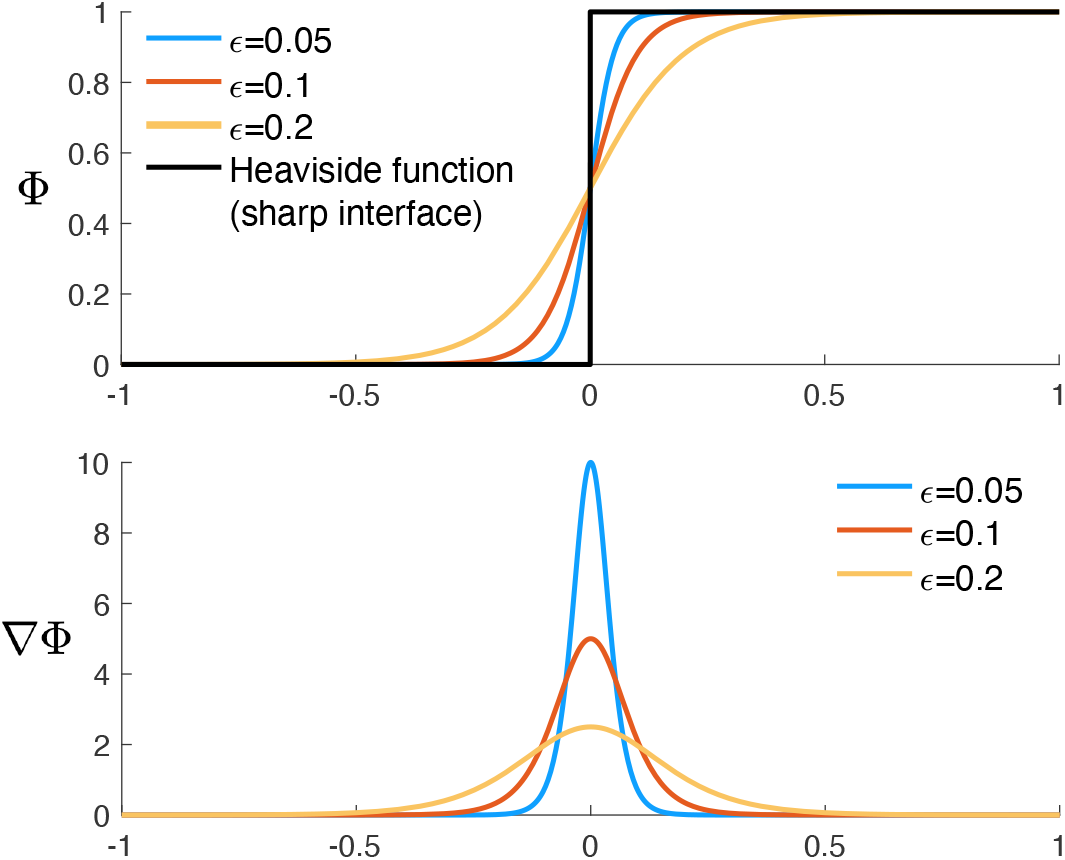
*Graphical representation of the diffuse interface approach in one dimension. Top: The phase-field function* Φ. *Bottom: The gradient of* Φ *used to approximate the location of the interface*.

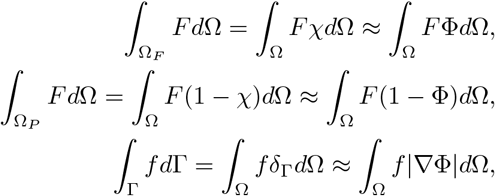

where *δ*_Γ_ is a Dirac distribution at the interface Γ.

Using the approximations above, and 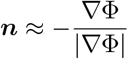, we can write the diffuse interface formulation of problem (2.8) as follows: Find (***v***, *p*_*P*_, ***η***, *p*_*F*_) ∈ *L*^2^(0, *T* ; *𝒱*_*F,D*_(Ω) × *H*^1^(Ω)) × *L*^∞^(0, *T* ; *𝒱*_*E*_(Ω)) × *H*^−1^(0, *T* ; *L*^2^(Ω)) such that for every (***w***, *ψ*_*P*_, ***ζ***, *ψ*_*F*_) ∈ *𝒱*_*F*_ (Ω) × *H*^1^(Ω) ×*𝒱*_*E*_(Ω) × *L*^2^(Ω), the following equality holds:

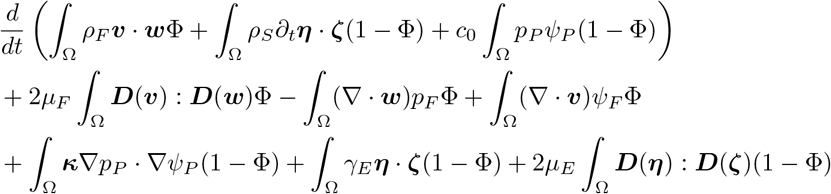

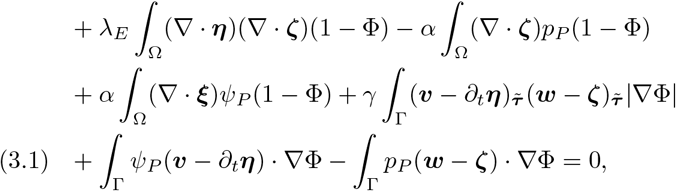

where we used the 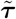 subscript to denote the “approximate” tangential component of a vector function at the diffused interface, defined, for a vector ***v***, as follows:

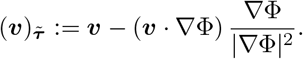

Tangent vectors that form a basis with ∇Φ can also be obtained directly from the phase-field function using the algorithm described in [21, 28].

We note that in order to write (3.1), the variables on each subdomain have to be extended onto the entire domain Ω. Since this procedure introduces singularities, we use the following regularization of Φ :

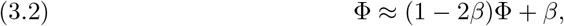

where *β* is a small positive number. Therefore, Φ ≥ *β* and 1 − Φ ≥ *β*. This regularization of Φ was used in the definition of problem (3.1). A phase-field method for the related Stokes-Darcy problems have been analyzed by the authors in [7] where existence of a weak solution was proved, and its convergence to the sharp interface solution was obtained.

Because the concentration equation (2.11) is already defined in Ω, we do not require regularization. In that case, *β* = 0. However, we use Φ to define the global velocity and diffusion:

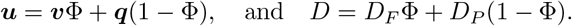

### Numerical discretization

Problems (3.1) and (2.11) are discretized in time using the Backward Euler method, and in space using a finite element method. Let *N* ∈ ℕ be the number of time steps, *T* the final time, and Δ*t* = *T/N* the time step. We define the discrete times *t*^*n*^ = *n*Δ*t*, for *n* = 0, …, *N* . We also denote by *h* a discretization parameter associated with the triangulation *𝒯* _*h*_(Ω) of Ω. For each *h*, we choose finite dimensional subspaces 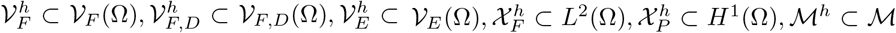 and 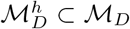 over the triangulation *𝒯*_*h*_(Ω). We use MINI elements [2] to approximate the fluid velocity and pressure, *𝒫*_1_ elements to approximate the Darcy pressure and the displacement, and *𝒫*_2_ elements to approximate the concentration.

The fully discrete fluid-poroelastic structure interaction problem is given as follows: given 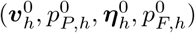, for *n* = 0, …, *N* − 1, find 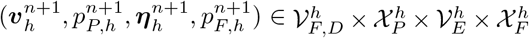 such that for every 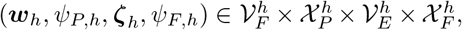, the following equality holds:

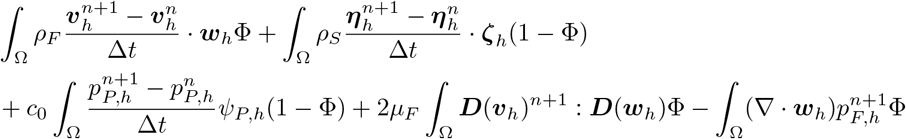

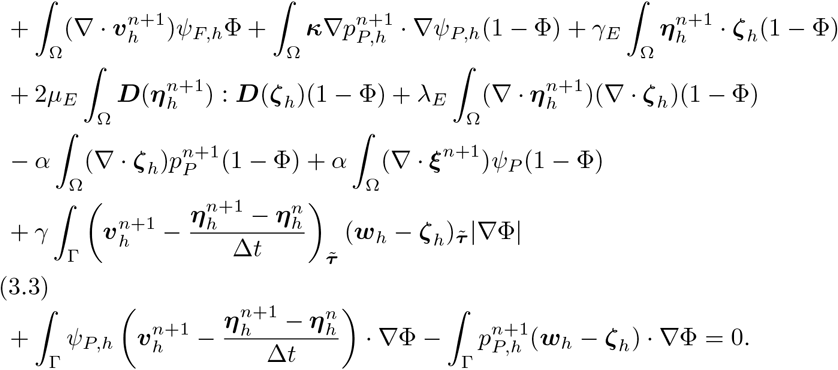

The fully discrete transport problem reads as follows: given *c*_0_, for *n* = 0, …, *N* − 1, find 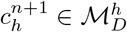 such that for every *s*_*h*_ ∈ *ℳ*^*h*^, the following equality holds:

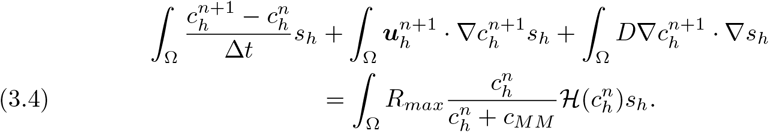

## 4. Computational Setting, Parameters Identification and Validation

In this work we investigate the fluid velocity, pressure and oxygen concentration in the three geometries/scaffold architectures shown in Figure 4. The first geometry consists of vertical ultrafiltrate channels drilled through a hydrogel, which is the simplest, and a standard procedure used in the design of bioartificial pancreas [26]. The second geometry consists of the branching channels, see the middle panel in Figure 4. This was inspired by the architecture of the branching vessels in the human body. The third geometry consists of the hexagonal channels filling the space between the inlet and outlet horizontal channels, shown at the bottom panel in Figure 4. This geometry was inspired by the biological (epithelial) tissues in which interstitial fluid flows through a network of irregularly arranged interstices between hexagonally shaped cells, which supports their structural and functional integrity [10]. To work with comparable fluid flow scenarios, the three geometries shown in Figure 4 were generated so that the total fluid channels’ area is the same in all three geometries.

**Fig. 4.**
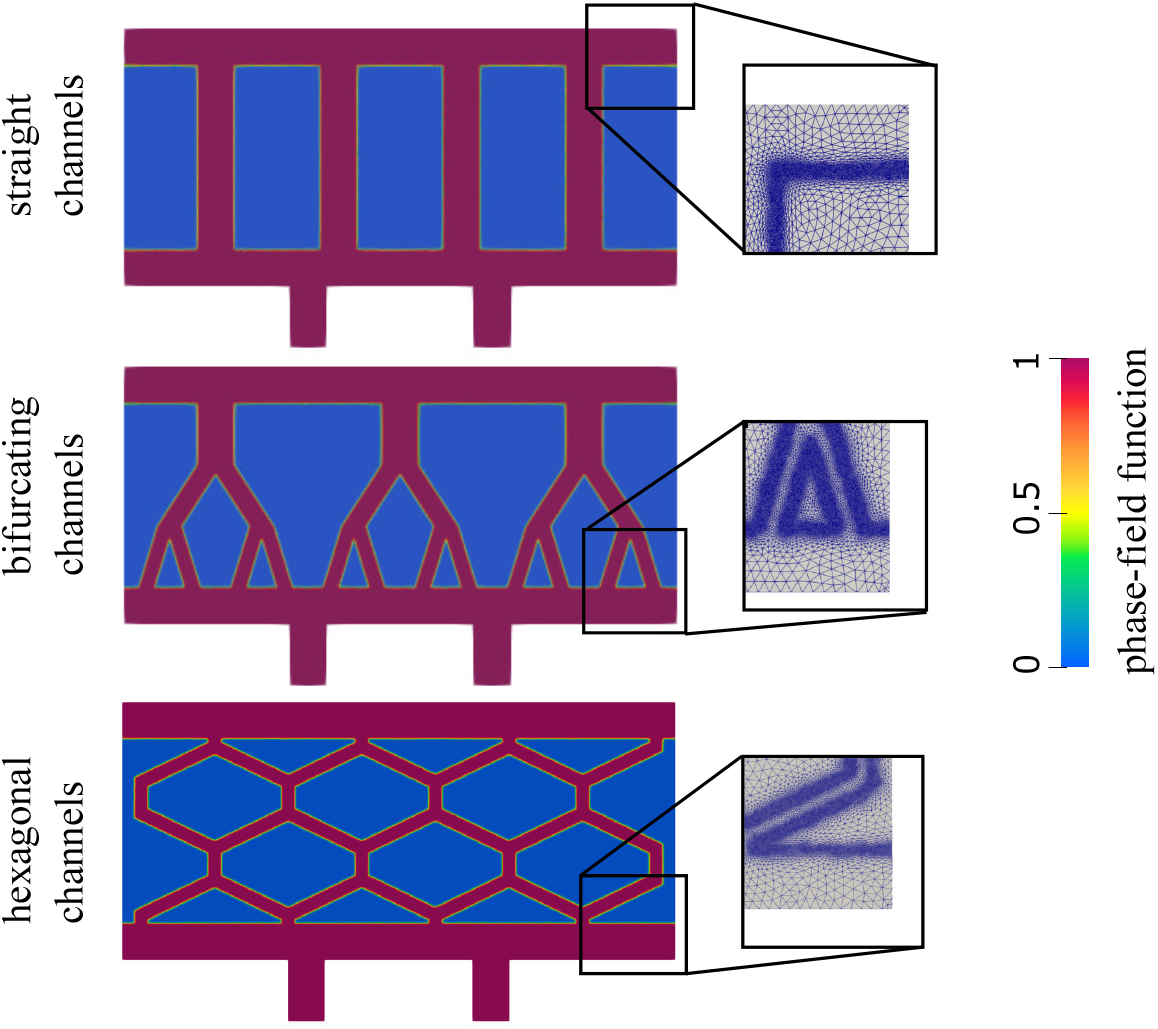
*The phase-field function for three different network configurations considered in this work. The zoom-in inserts show the computational mesh. The mesh is refined in the areas with large* |∇Φ|, *which approximate the interface*.

### 4.1 Computational settings

The size of the computational domain is 0.9 × 0.42 cm, corresponding to the entire scaffold Ω. Two outlet channels are added on the bottom (see Figure 2) to simulate the actual outlet channels shown in Figure 1. Each outlet channel has a height of 0.1 cm and length of 0.06 cm. In each of the three different geometries/architectures of the scaffold Ω, there is a top and a bottom horizontal channel of width 0.06 cm. The same width of 0.6 cm is used for the vertical channels in the first, top configuration. The size of the channels in other configurations is chosen so that the total channel area equals the channel area of the first configuration. In other words, the channels occupy the same area in all three configurations. To computationally capture different geometries/architectures of thin channels distributed within Ω we use a phase-field function. The phase-field function for each of the three geometries is shown in Figure 4. Here, red color corresponds to the value of the phase-field function Φ equal to one, and blue corresponds to Φ = 0. The computational mesh is refined around the interface between the red and blue regions where the gradient of the phase-field function | ∇ Φ | is large. This is done as follows. We first set Φ = 1 in the region defined by the channels and zero elsewhere. Then, we adapt the mesh and redefine Φ on the finer mesh. After that, we solve the following Allen-Cahn problem to allow a smooth transition between 1 and 0:

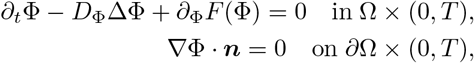

where *D*_Φ_ = 0.01 and 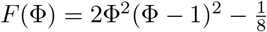 is the standard form of a double-well potential [15]. Finally, the procedure of adapting the mesh, redefining Φ and solving the Allen-Cahn problem is repeated one more time. The resulting computational mesh is shown in the zoom-in inserts in Figure 4 (right).

### 4.2 Parameter identification

The fluid in Ω_*F*_ represents the filtered blood plasma, which enters the bioartificial organ Ω from an artery via an anastomosis graft, not shown in Figure 4. In encapsulated organs, the blood from the anastomosis graft is filtered through semipermeable nano-pore membranes, and the filtered blood plasma enters a gasket from which the flow of plasma filtrates through the biocompatible hydrogel Ω_*P*_ toward the cells. The horizontal channels represent the gasket containing the blood plasma, and 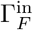 corresponds to the location of the semipermeble membranes through which blood plasma enters the top gasket. At 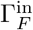 we prescribe the inlet velocity, which is taken to be ***v***_*in*_ = 3.5 cm/s. This value agrees with the experimentally derived relationship between the flow and pressure gradient through nanoporous membranes reported in [16]. It also agrees with the data used in threedimensional simulations of flow through an entire bioartificial pancreas, studied in [31].

The parameters used in the blood plasma simulations are standard: the kinematic viscosity is set to *v* =1 cm^2^/s and density (per 2*D* unit area) is *ρ*_*F*_ = 0.04 g/cm^2^.

The parameters for the poroelastic structure describing the hydrogen scaffold can be obtained from [31, 14]. We took the 2*D* poroelastic matrix density to be *ρ*_*S*_ = 1.2 g/cm^2^, and the Young’s modulus of 2.5 · 10^5^ dyne/cm. The Poisson’s ratio is set to 0.49 and the spring constant is *γ*_*E*_ = 10^6^ dyne/cm^3^. The pressure storage coefficient is taken as 10^−7^ cm/dyne, and the permeability is *κ* = 10^−5^ cm^2^ s/g. The slip constant is *γ* = 10^3^ g/(cm s). We set the Biot-Willis parameter *α* to 1. See [31] for the Biot parameters.

Oxygen transport in the human vascular system and tissues has been studied by many authors [4, 5, 17, 23, 22, 25]. We adopt the approach from [4] to study oxygen transport in the gasket with oxygen diffusion coefficient given by *D*_*F*_ = 3 · 10^−5^ cm^2^/s [4]. In the agarose gel, the oxygen diffusion coefficient used in our simulations is *D*_*P*_ = 1.3 · 10^−5^ cm^2^/s, which is the value that was estimated in rat pancreatic islets and reported in [3, 25]. The maximum oxygen consumption rate is *R*_*max*_ = − 3.4 · 10^−8^ mol/cm^2^ s, the Michaelis-Menten constant is *c*_*MM*_ = 10^−9^ mol/cm^2^ and the critical oxygen concentration is *c*_*CR*_ = 10^−10^ mol/cm^2^, all obtained from [5]. The concentration of oxygen at the fluid inlet is *c*_*in*_ = 2 · 10^−7^ mol/cm^2^ [11]. All units are adapted to the 2*D* case.

We first perform simulations of the fluid-poroelastic structure interaction problem using Δ*t* = 10^−3^ and *T* = 1 s, which is when a steady state is reached. Using the velocities at the final time of the simulation, we then solve the advection-reaction-diffusions problem with Δ*t* = 5 ·10^−2^ and *T* = 200. All computations are performed within the platform of the finite element software FreeFem++ [18].

To trust the simulations obtained using our diffuse interface method, we validate our computational solver by comparing the results of the diffuse interface method with the results obtained using the “classical” sharp interface approach, on a simpler geometry, as we discuss next. Recall that the main reason for not using the sharp interface approach is the difficulty in generating new scaffold geometries for each new test case. This will be particularly important in our next research phase in which a geometric optimization mathematical model and computational solver will be developed to study optimal design of channels’ distribution in bioartificial organ scaffolds for advection-enhanced oxygen and nutrients supply to the transplanted cells.

### 4.3 Numerical method validation

To validate our diffuse interface solver, we focus on a specific problem characterized by a domain geometry comprising a main channel branching into two, each of which further bifurcates into additional two channels. See Figure 5. We solve the fluid-poroelastic structure interaction problem and the advection-reaction-diffusion problem using a sharp interface approach. The sharp interface solver is based on a classical, monolithic, fluid-poroelastic structure interaction approach. The solver was developed and validated in [8, 24]. In particular, the Backward Euler method is used to discretize the problem in time, and a finite element method is used for spatial discretization. The same finite element spaces are used for the diffuse and sharp interface methods. The results of the sharp interface solver are then compared to the results obtained using our diffuse interface solver discussed in this manuscript. All the parameter settings are the same as the ones described in Section 4. Since the permeability is small, a finer mesh close to the interface is required for both sharp and diffuse interface solvers. We use the same mesh in both cases, consisting of 40,239 points and 80,326 elements. A comparison of the Stokes and Darcy pressure and velocity magnitude is shown in Figure 5. Slight differences are observed in the pressure, however, an excellent agreement is obtained for the velocity. In the right panel of Figure 5 we show a comparison of the concentrations at *T* = 200 obtained using the sharp interface model and the diffuse interface model. Excellent agreement is observed.

**Fig. 5.**
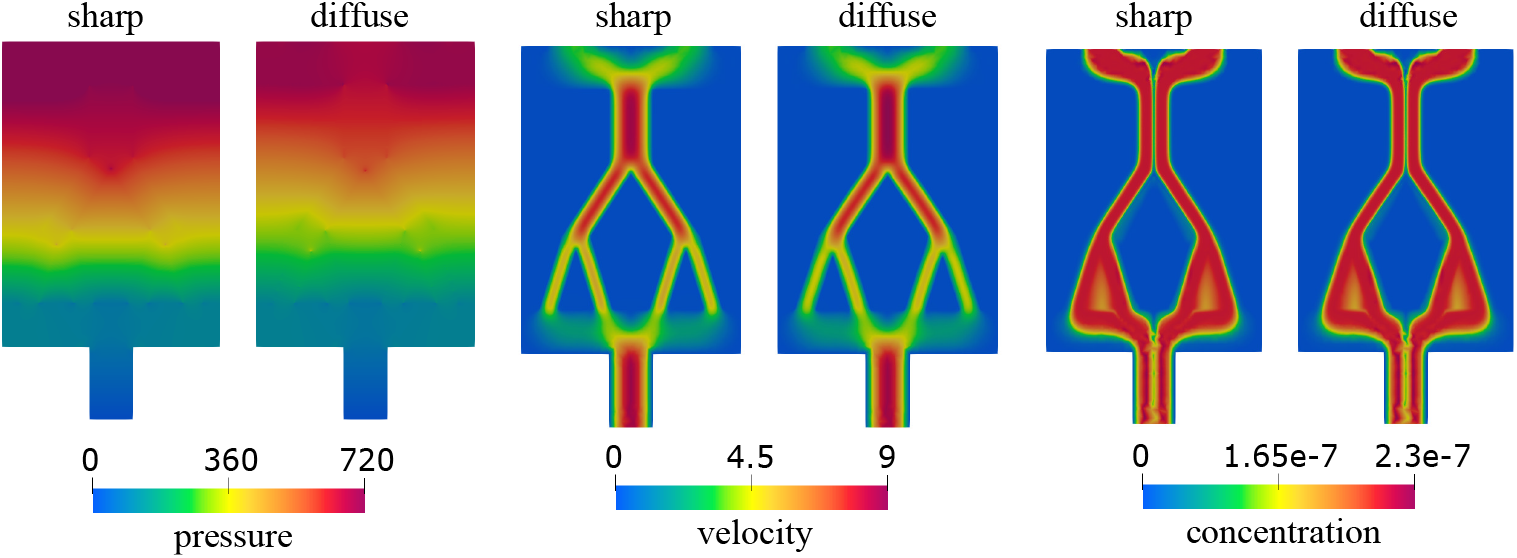
*A steady-state solution for the Stokes and Darcy pressure (left) and velocity magnitude (middle) obtained using the sharp interface model and the diffuse interface model. The right-most panel shows concentration obtained at T* = 200.

Encouraged by these results, we use the diffuse interface method described above to study the flow and concentration of oxygen in the geometries shown in Figure 4. This is presented next.

## 5. Numerical results

As mentioned earlier, our goal is to study blood plasma flow and oxygen concentration in three different scaffold architectures, as described in Section 4. We are interested in a scaffold architecture that provides concentration of oxygen that is as uniform throughout the scaffold as possible and above the minimum concentration for uninhibited maximal insulin production *c*_*opt*_ = 5 · 10^−8^ mol/cm^2^, given in [25]. To do this, we use the diffuse interface method to simulate fluid-poroelastic structure interaction providing advection velocity of blood plasma carrying oxygen, and we use the advection-reaction-diffusion solver to calculate oxygen concentration in the entire scaffold, utilizing the advection velocity from the fluid-poroelastic structure interaction simuations. Both models are described in Section 2. The results of the simulations are shown in Figures 6 and 7 below. More precisely, in Figure 6 we plot the total fluid velocity, defined as

**Fig. 6.**
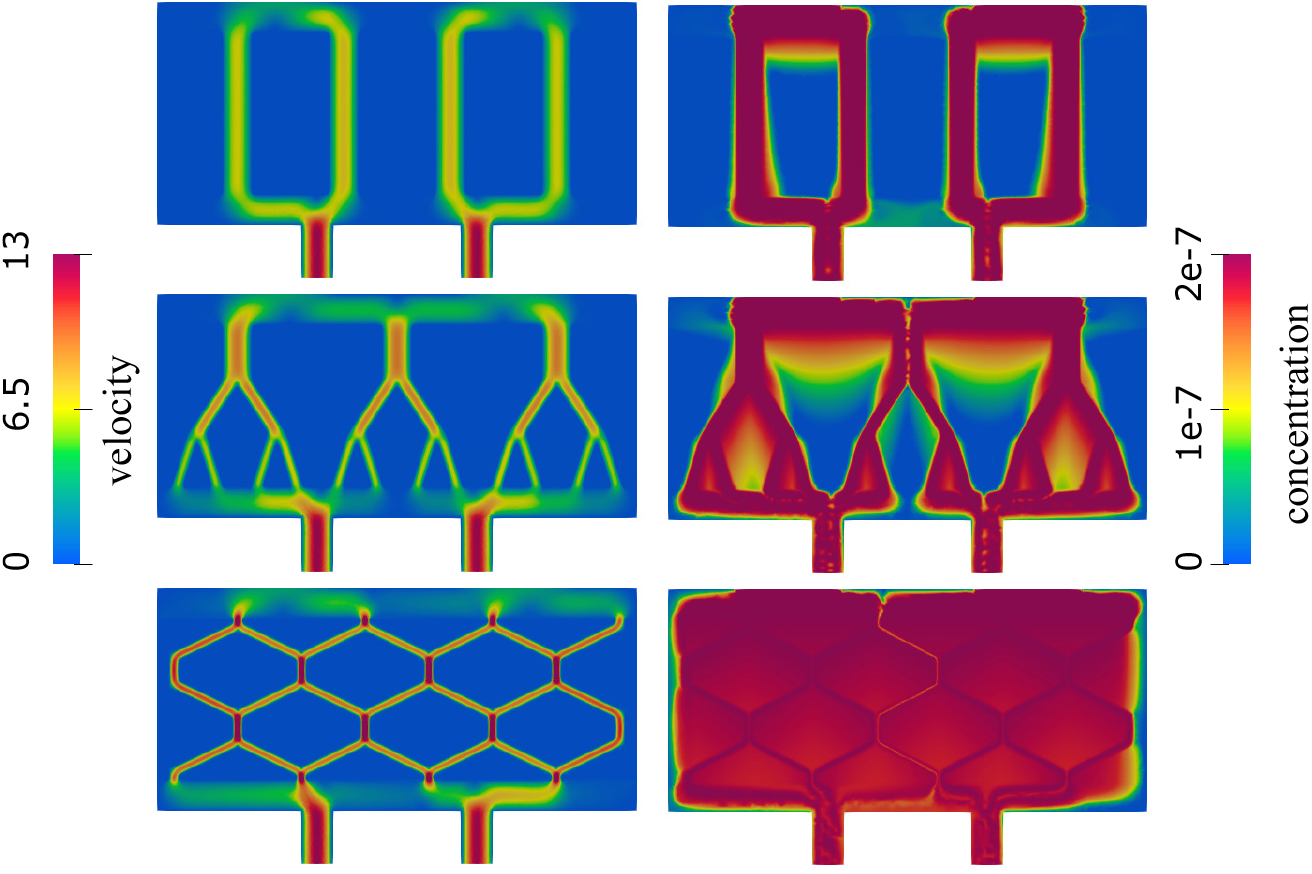
Total velocity magnitude (left) and concentration (right) in a network consisting of straight channels (top), bifurcating channels (middle) and a hexagonal network of channels (bottom).

**Fig. 7.**
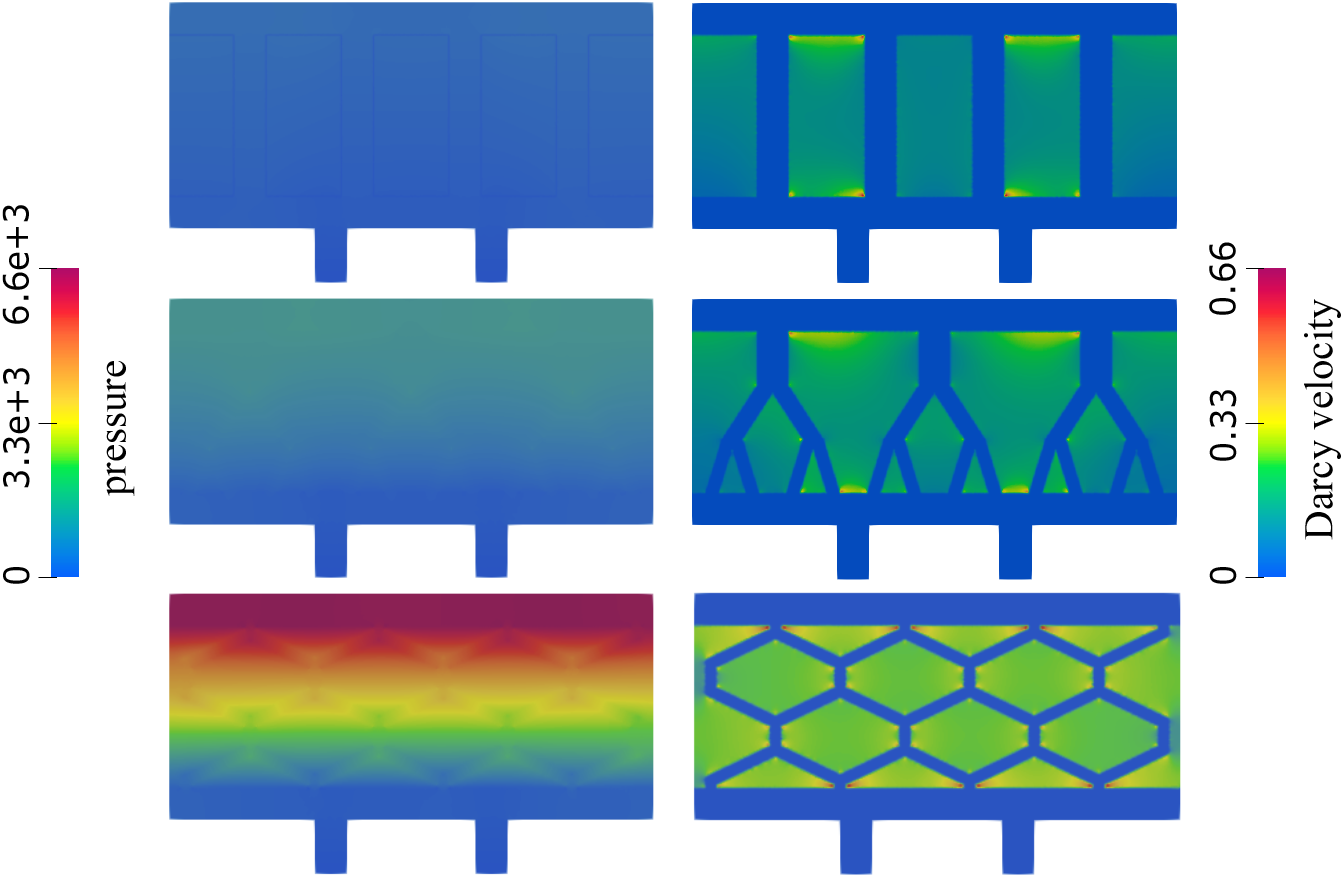
Pressure (left) and the Darcy velocity (right) in a network consisting of straight channels (top), bifurcating channels (middle) and a hexagonal network of channels (bottom).

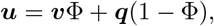

and oxygen concentration for the three different scenarios.

We observe in Figure 6 that in the case of vertically drilled channels, most of the flow is through the vertical channels, with Darcy velocity, shown in Figure 7, highest in the top part of the poroelastic hydrogel. This correlates with the values of oxygen concentration, shown in Figure 6 right, which are highest in the top part of the poroelastic hydrogel, and in the channels (as expected). There are large regions in between the channels where oxygen concentration is below the critical value *c*_*opt*_ = 5 ·10^−8^ mol/cm^2^ [25], indicating regions of impaired insulin production.

In the case of bifurcating channel network, shown in the middle panels on Figures 6 and 7, we see higher velocity values in the top two generations of branching channels (parent and daughter channels), and larger regions of higher Darcy velocity, shown in light green and yellow, in between the branching vessels. Consequently, we observe larger regions of higher oxygen concentration in between the channels than in the case of vertical channels, see Figure 6 right, middle panel. However, there are still large regions in between the branching trees that have low levels of oxygen concentration, shown in blue, where insulin production is inhibited.

Finally, in the case of the hexagonal channel network, shown in the bottom panels of Figures 6 and 7, the channels have the smallest radius and the fluid velocity through the hexagonal network is the highest. Darcy velocity in this case is nontrivial in the entire region corresponding to the poroelastic hydrogel. As a result, oxygen concentration is high in the entire hydrogel region, and it is very close to the uniform concentration above the critical value *c*_*opt*_, as desired, providing oxygen supply to the cells in the entire hydrogel that is above the critical value above which normal insulin production takes place.

### Two additional geometries

We consider two additional geometries to gain a better understanding of why the hexagonal geometry is associated with improved oxygen concentration levels. Since hexagonal channel architecture described above has narrower channels that are at an angle with respect to the pressure drop-driven dominant vertical flow, we separate the influence of the two geometric factors and consider the following two geometries: one consisting of narrow vertical channels with a small radius (the width of each channel is equal to one third of the width of vertical channels used above, shown in Figure 4), and the other consisting of the zig-zag (angular) channels with the radius determined by the constraint that the total channel area is the same as in the vertical channel case. See Figure 8.

**Fig. 8.**
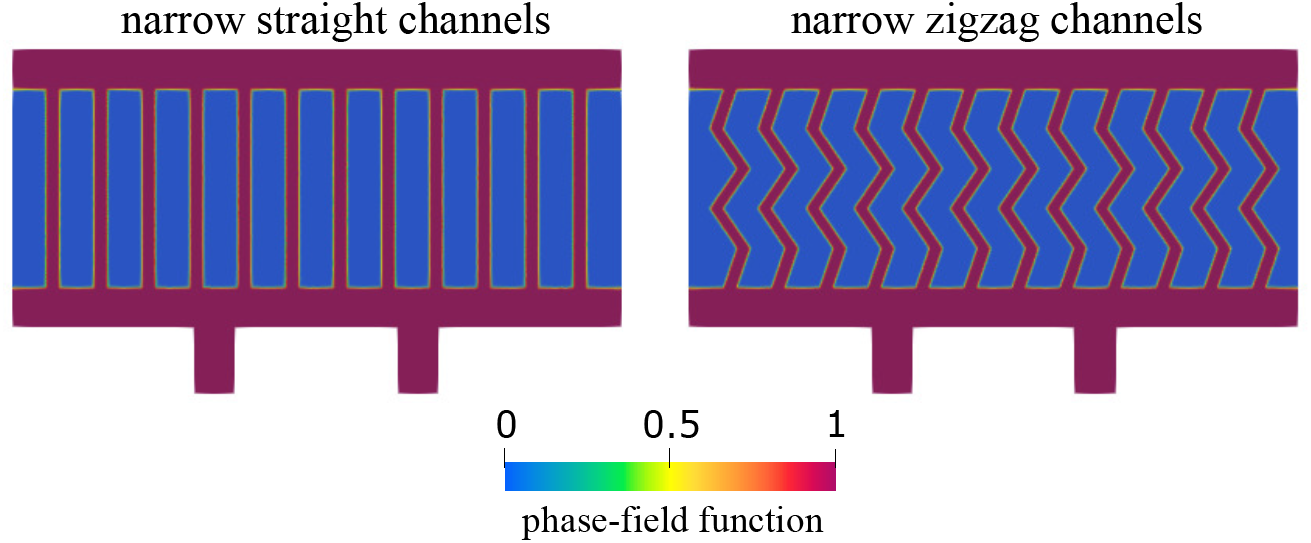
Geometries consisting of narrow straight channels (left) and narrow zigzag channels (right).

The flow velocity and concentration obtained in the new geometries are shown in Figure 9. In both geometries, the peak velocity in the channels is equal to 13 cm/s. However, significant differences can be seen in concentration. We note that having narrow channels improves the transport through the poroelastic medium compared to having fewer wider channels (see top-right panel in Figure 6). However, having a network with channels at an angle with respect to dominant flow leads to increased oxygen levels overall, especially at places where the flow changes direction. An explanation for this observation is the fact that flow through porous interfaces is largest when the angle between flow direction and the interface is large.

**Fig. 9.**
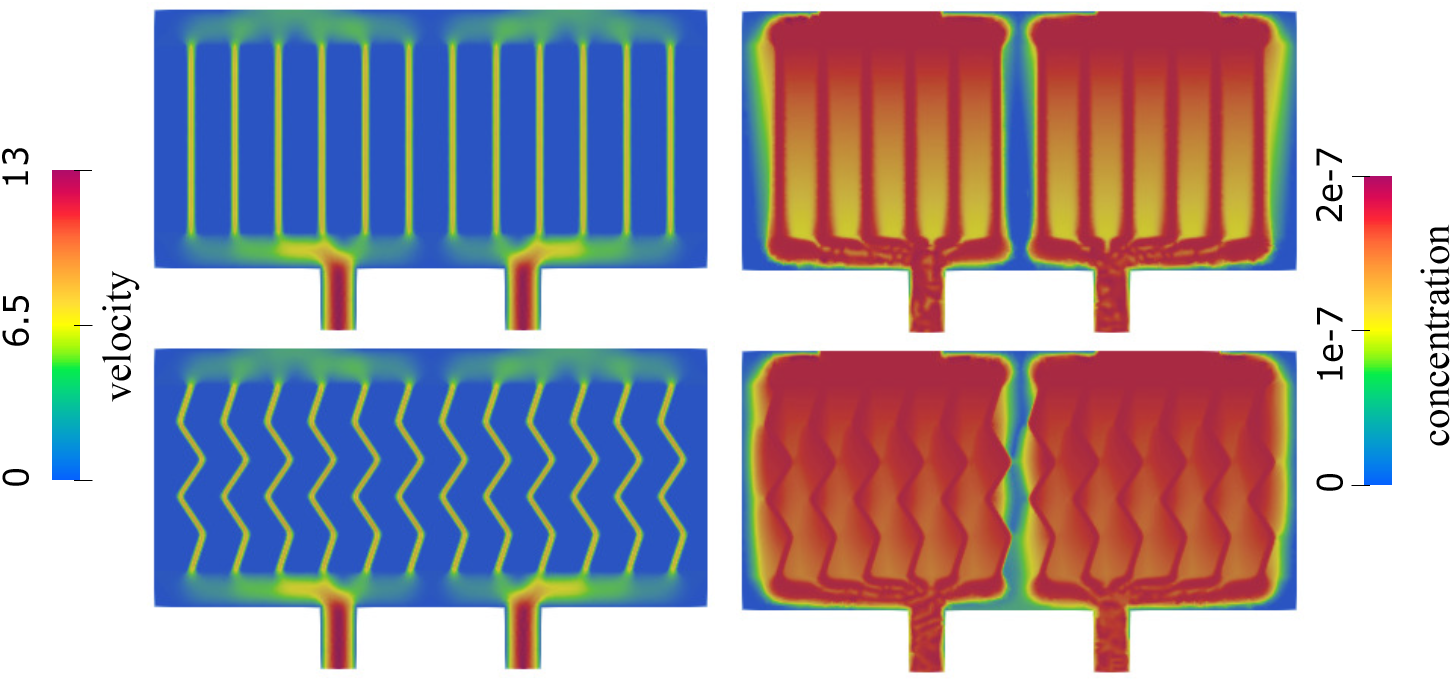
Total velocity magnitude (left) and concentration (right) in a network consisting of narrow straight channels (top) and narrow zigzag channels (bottom).

This is evident in the right panel of Figure 10 where Darcy velocity is shown. The peak Darcy velocity is twice as large when zigzag channels are used compared with the straight channels. To improve the visualization of the peak velocity area, we show the flow in two geometries on different scales. When straight channels are used, the flow is largest closest to the top and bottom channels. In zigzag channels, the flow is largest in the middle of the domain. This is an interesting observation that significantly improves oxygen supply to the cells located in the middle of the proelastic hydrogel. We also show the total pressure in the two geometries in Figure 10. Our results show a significantly larger internal scaffold pressure in case of the zigzag network, which is associated with increased Darcy flow through the hydrogel.

**Fig. 10.**
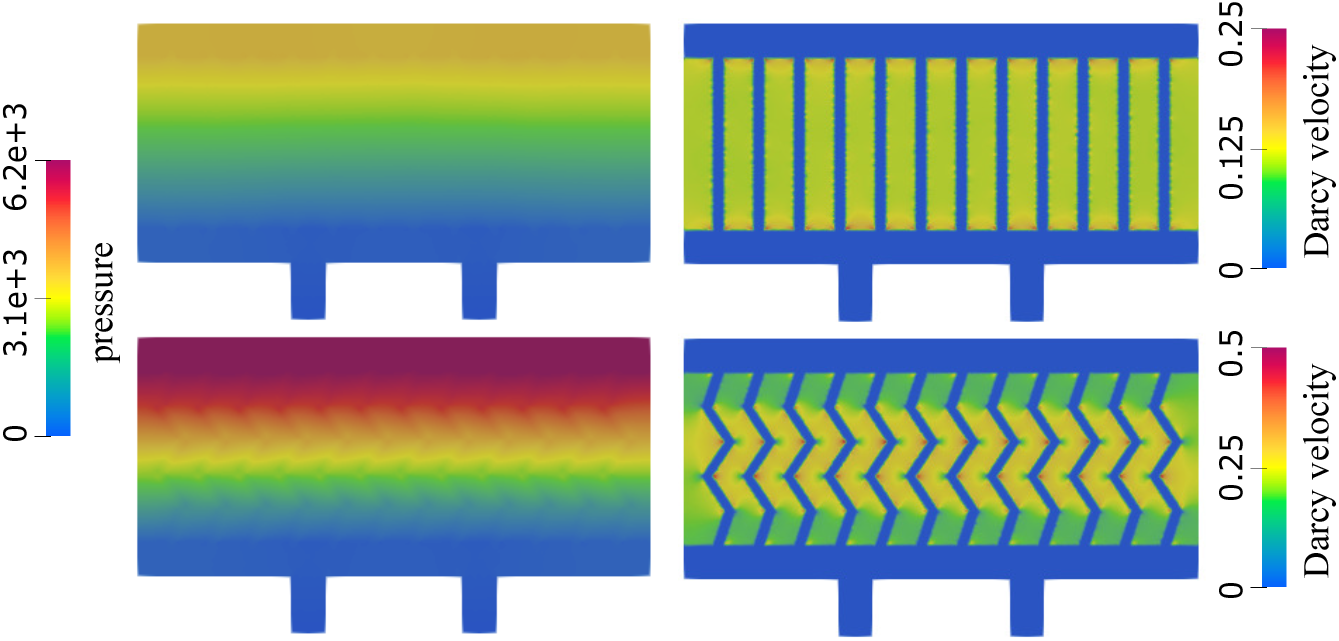
*Pressure (left) and the Darcy velocity (right) in a network consisting of narrow straight channels (top) and narrow zigzag channels (bottom)*.

### Uninhibited maximal insulin production regions

Finally, we further quantify the behavior of the three scaffold architectures shown in Figure 4 by considering the performance of the three geometries in terms of the uninhibited maximal insulin production by the transplanted *β*-cells [25]. Namely, motivated by the results in [25] we use a threshold for uninhibited maximal insulin production by the transplanted islets of *c*_*opt*_ = 5 ·10^−8^ mol/cm^2^ to identify the regions within tissue in which the islet function is compromised. More precisely, we investigate the regions within each scaffold in which the oxygen concentration is higher than the threshold for uninhibited maximal insulin production, *c*_*opt*_, and compare the areas of those regions that support islet function. This area is computed only in the poroelastic domain using the following expression:

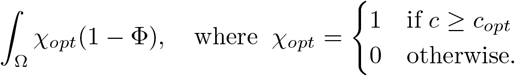

The regions are visualized in Figure 11. Blue color shows the regions of uninhibited insulin production, while the insulin production is compromised in the regions shown in red. The results are quantified in Table 1. As expected, the hydrogel geometry with the smallest area of oxygen levels above *c*_*opt*_ is obtained for the network consisting of straight channels (13% of total area), followed by the bifurcating channels, which have the uninhibited maximal insulin production area equal to 51.74% of the total area. Finally, the most efficient insulin production is observed in the hexagonal geometry, where more than 97% of the poroelastic region is above the uninhibited maximal insulin production threshold *c*_*opt*_.

**Table 1.** The area where the concentration is larger than the oxygen threshold of uninhibited maximal insulin production for all the geometries considered in this manuscript.

| Geometry | Total area (cm<sup>2</sup>) | Relative area |
| --- | --- | --- |
| straight network | 0.026 | 13.23% |
| bifurcating network | 0.102 | 51.74% |
| hexagonal network | 0.192 | 97.13 % |
| narrow straight network | 0.172 | 86.94 % |
| narrow zigzag network | 0.187 | 94.24 % |

**Fig. 11.**
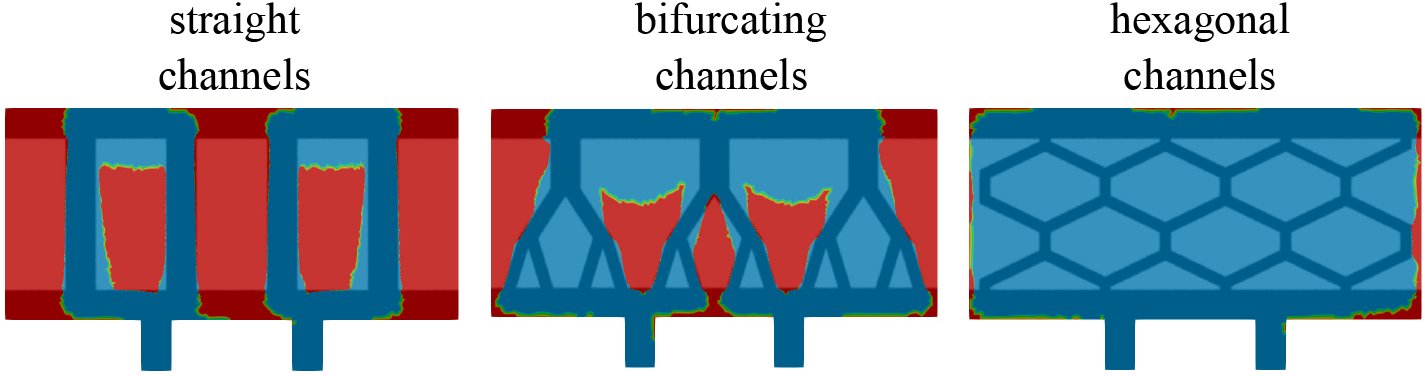
Function χ_opt_ superimposed with the phase-field function indicating the channel geometry. Regions in red show areas where the insulin production is inhabited, while regions in blue show areas of uninhabited insulin production.

## 6. Three-dimensional simulations

We conclude this manuscript by demonstrating that the 2D results presented above accurately represent real-life 3D scenarios. In particular, we focus on a 3D scaffold consisting of hexagonal channels, as shown in Figure 12 below. Each vertical slice of this geometry is identical to the 2D geometry shown at the bottom of Figure 4. The dimensions of the 3D spatial domain are 0.9 × 0.42 × 0.3 cm. Here 0.3 cm is the added thickness of the 3*D* domain. We maintained the same inlet and outlet setup as in the 2D case, and also kept all other parameters unchanged.

**Fig. 12.**
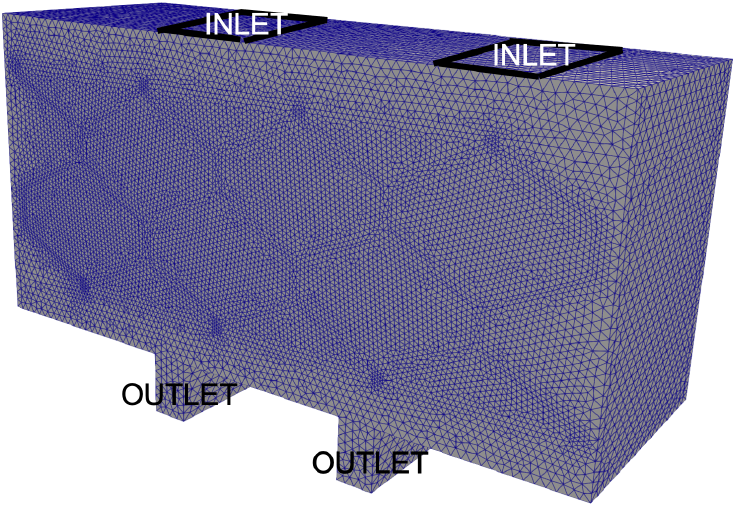
Computational Domain: The computational domain was discretized using conformal tetrahedral elements, resulting in a total of 926K elements.

For the boundary conditions on the front and back wall of the computational domain we imposed the no-slip boundary condition for the free fluid velocity modeled by the time-dependent Stokes equations, and zero normal flux for Darcy velocity for the Biot equations. Displacement of the poroelastic matrix was also set to be equal to zero on the front and back walls of the chamber. On the rest of the boundary, we implemented the same boundary conditions as in the 2*D* case. Similarly, at the inlet, shown in Figure 12, Dirichlet boundary condition was imposed for the fluid velocity, and at the outlet, zero normal stress was imposed, as in the 2*D* case. The Dirichlet velocity imposed at the inlet is given by the uniform velocity profile with the magnitude of 3.5 cm/s, pointing downwards in the direction of the cell chamber.

A monolithic solver reported in [8, 24, 30] was used to solve the 3D linearly coupled Stokes-Biot problem. Figure 12 depicts the computational domain, which was discretized using conformal tetrahedron elements. Taylor-Hood elements (P2-P1) were used for the free fluid velocity and pressure, P2 for the poroelastic structure displacement, P2 for filtration velocity and P1 for Darcy pressure. We set the time step to be Δ*t* = 0.005*s*, and allowed the simulation to run until it reached a steady state at *T* = 1s. As in the 2D case, the flow is driven by the pressure drop in the vertical direction generated by the inlet velocity and outlet normal stress data.

We examined the following quantities and compared them with the 2D simulation results: the free fluid velocity and pressure (at the inlet and outlet channels, and in the hexagonal channels throughout the poroelastic medium), the filtration velocity and Darcy pressure within the poroelastic structure, and the deformation of the poroelastic structure.

### Velocity

Figure 13 shows a comparison of the velocity obtained using 2D simulations (left) and 3D simulations (middle and right).

**Fig. 13.**
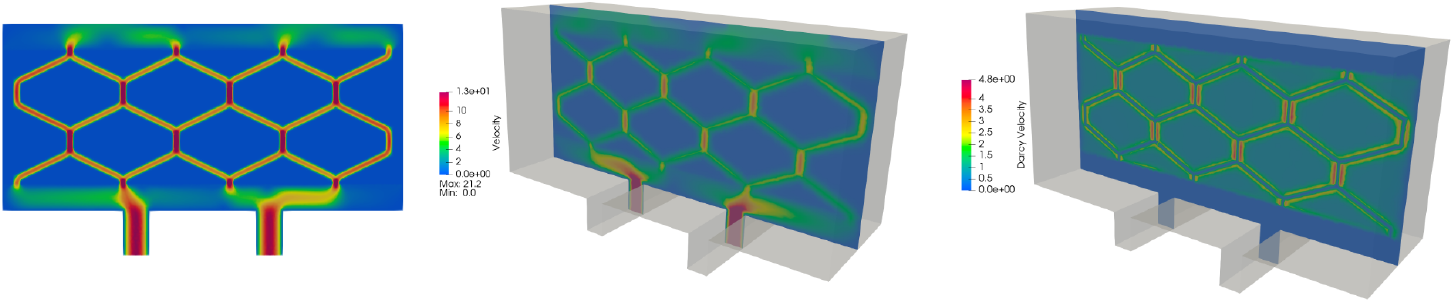
Velocity comparison. Left: 2D simulation showing free fluid and Darcy velocity ranging from 0 to 13 cm/s. Middle: 3D simulation showing velocity in the inlet, outlet and hexagonal channels ranging from 0 to 13 cm/s. Right: 3D simulation showing only Darcy velocity ranging from 0 to 4.8 cm/s.

The middle panel shows only the free fluid (channel) velocity, while the right panel shows only Darcy velocity (poroelastic medium). Notice two different scales for the middle and right panel. We see that in both 2D and 3D scenarios the fluid velocity ranges from 0 to 13 cm/s, and that the highest velocity is achieved in the vertical channels, as expected due to the dominant pressure drop in the vertical direction.

### Pressure

Next we compared the fluid pressure. Figure 14 shows a comparison of the pressure obtained using 2D simulations (left) and 3D simulations (middle and right). In all three panels the pressure ranges from 0 to around 6,000 dyn/cm^2^. The panel on the left shows the combined 2D free fluid and Darcy pressure, while the figures on the right show separate 3D Darcy pressure (middle) and 3D free fluid pressure (right). When combined, the middle and right panels are in good agreement with the panel on the left, indicating that the pressure distributing obtained using 2D simulations is indeed, very close to the pressure distribution obtained using 3D simulations.

**Fig. 14.**
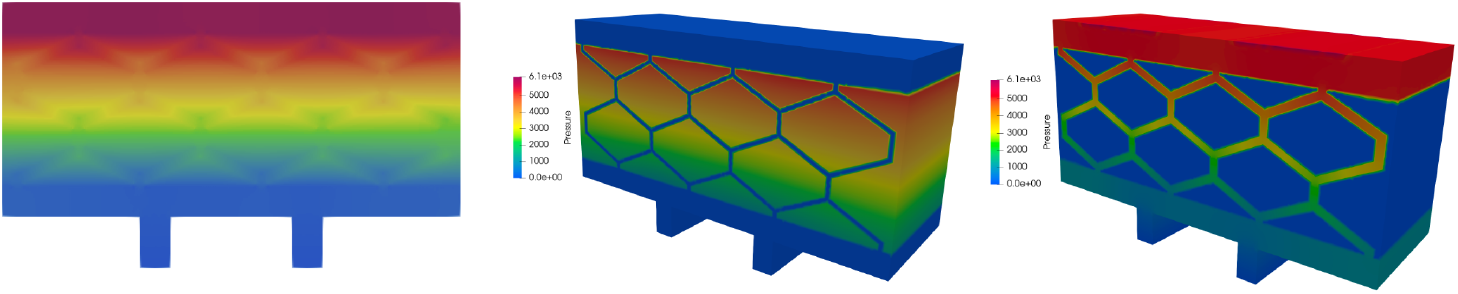
Pressure comparison. Left: 2D simulation showing free fluid and Darcy pressure. Middle: 3D simulation showing Darcy pressure. Right: 3D simulation showing pressure in the inlet, outlet and hexagonal channels. The pressure scale is the same in all three panels – from 0 to around 6,000 dyn/cm^2^.

### Streamlines

To further compare 2*D* versus 3*D* spatial effects on the solution, we investigated the fluid velocity streamlines inside the entire 3*D* scaffold. This is shown in Figure 15. We can see that 2*D* effects are dominant over 3*D* effects, since the streamlines appear to be largely parallel to each other in the direction perpendicular to the plane containing the hexagonal channels, indicating that 2*D* simulations approximate well the leading features of the fluid flow in the scaffold.

**Fig. 15.**
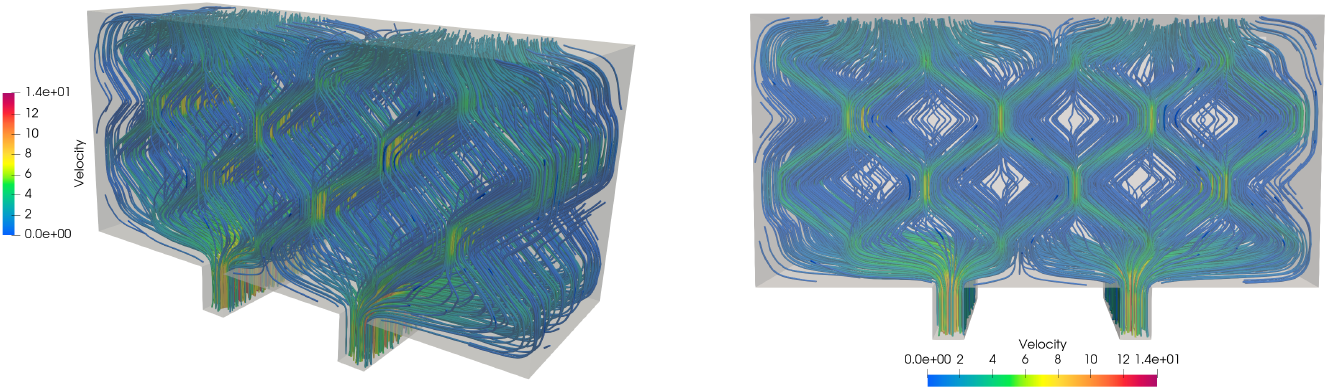
Streamlines generated by the fluid and filtration velocity within the scaffold. The color of the streamlines corresponds to the velocity magnitude. **Left**: Side view, **right**: Front view.

### Displacement

Using 3*D* simulations we also investigated the total displacement of the poroelastic matrix as time increases toward the time at which the steady state solution is achieved. The magnitude of displacement at the steady state is shown in Figure 16 (left) and the vectors showing the displacement vector field are shown in Figure 16 (right). We observe that all cell compartments have expanded from their original shape, with the compartments closest the the inlet experiencing larger expansion than those closest to the outlet. We argue that this expansion is a result of flow saturation inside the cell compartments. This also results in a constriction of the original hexagonal flow channels, potentially increasing the proportion of flow passing through the cell compartments.

**Fig. 16.**
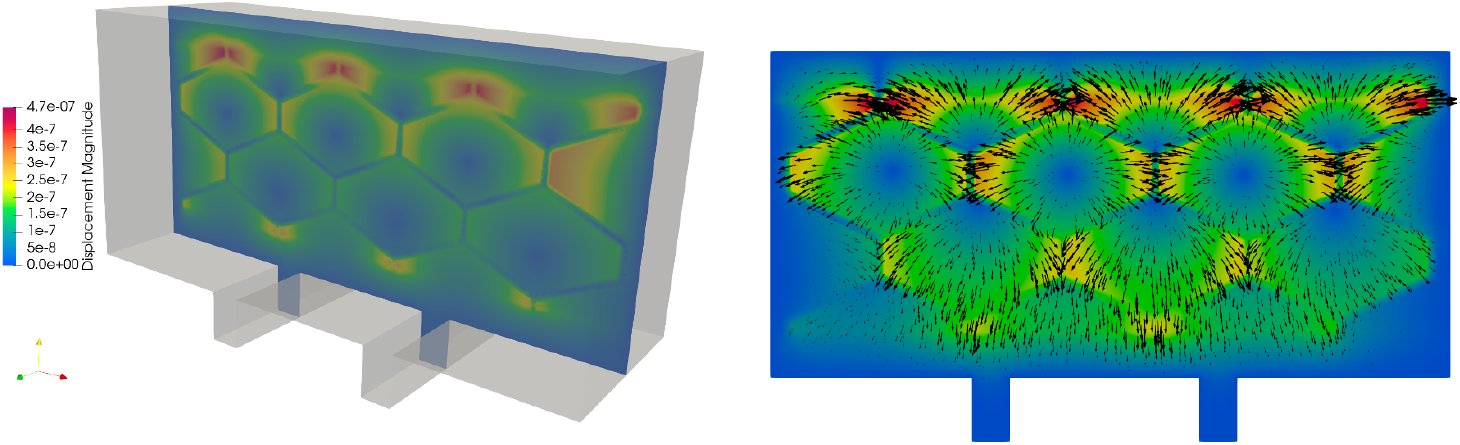
Displacement of the cell chamber at the steady state. Left: Magnitude of displacement. Right: Displacement vector field.

### Oxygen concentration

Using 3*D* simulations we also investigated oxygen transport within the scaffold by solving the advection-reaction-diffusion problem described in Section 2.2. The advection-reaction-diffusion problem was solved with the time step of Δ*t* = 1*e* − 5*s*. In Figure 17 we illustrate the evolution of oxygen concentration from *T* = 0 to *T* = 0.15 seconds and then give the final, steady state solution oxygen concentration achieved at *T* = 1s.

**Fig. 17.**
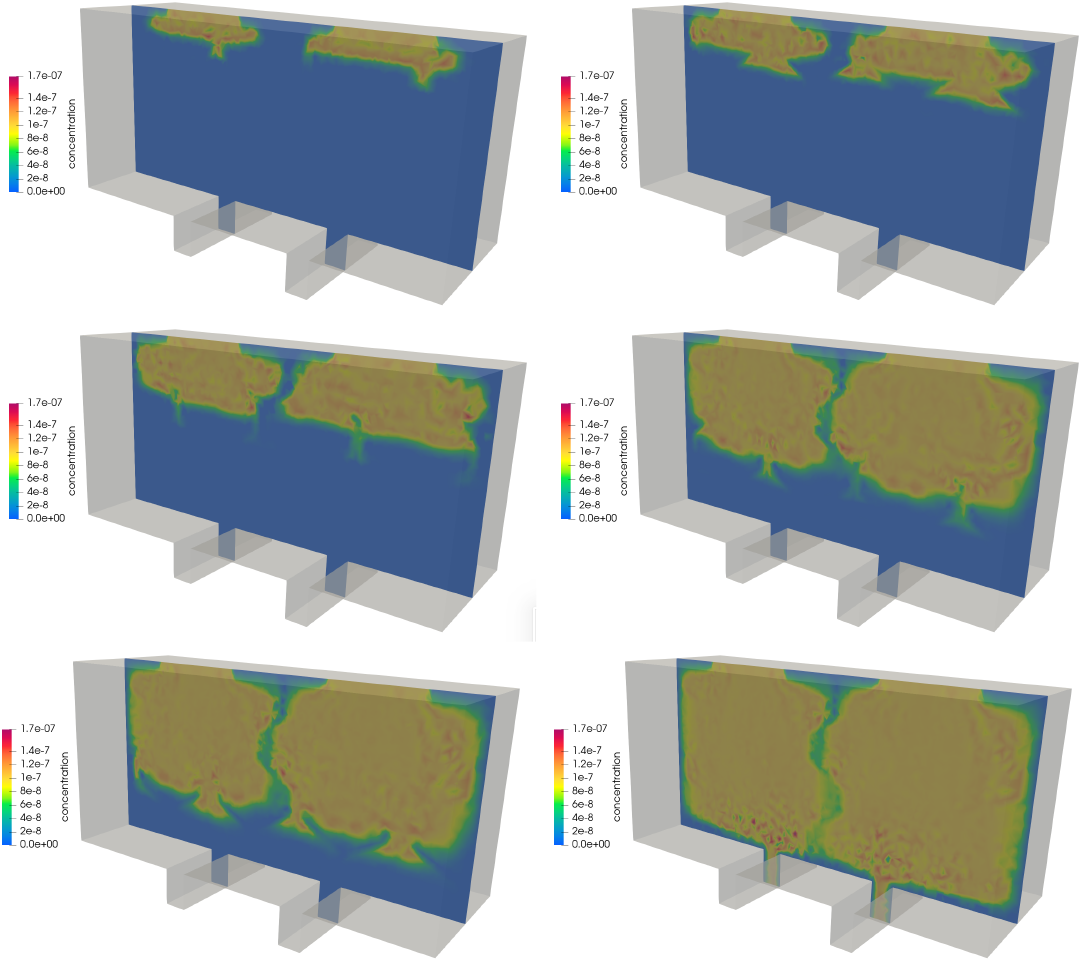
A series of screenshots depicting the transport of oxygen concentration from t = 0 s to t = 1 s corresponding to the steady state solution.

Figure 17 shows that indeed, the final oxygen concentration at *T* = 1 is nearly uniformly distributed throughout the chamber, except for the thin vertical region roughly half way between the two inlets. This is similar to the result obtained using 2*D* simulations, depicted in Figure 6 bottom right, which shows nearly uniform distribution of oxygen concentration except for a thin line located roughly in the middle between the two inlets. We conclude that, again, 2*D* simulations provide a good insight into oxygen concentration distribution within the cell chamber. Additionally, this result indicates that placing the two inlets closer to each other might improve uniform distribution of oxygen concentration within the chamber to include the central region between the two inlets.

## 7. Conclusions

We devised a computational approach within the diffuse interface framework to explore the influence of scaffold architecture geometry on oxygen transport within biological scaffolds commonly employed in bioartificial organ engineering, with a specific focus on the bioartificial pancreas. To achieve this objective, we introduced a multi-physics model comprising a fluid flow component and an advection-reaction-diffusion model to analyze oxygen concentration within the scaffold. The fluid flow model incorporates the time-dependent Stokes equations coupled with the Biot equations, characterizing the behavior of a poroelastic medium representing the poroelastic hydrogel used in the scaffold design.

We explored three biologically inspired scaffold architecture geometries: vertically drilled channels, branching channels, and a hexagonal channel network. From a computational standpoint, one of the primary challenges in addressing problems with multiple varying geometries lies in generating new computational geometries and designing appropriate matrices for spatial discretization. These matrices describe the unknown variables, such as fluid velocity, pressure, and oxygen concentration, across different computational domains. To streamline mesh generation in complex geometries, we introduced a diffuse interface approach in this study. In the diffuse interface approach, the unknown variables are defined across the entire scaffold domain, with the specific geometry of the channel network captured by redefining only the phase-field function. This simplification proves crucial not only for our current work but also for future endeavors, where we aim to develop a geometric optimization solver. This solver will simplify the generation of numerous channel geometries to optimize scaffold architecture. It is important to notice that a drawback of the diffuse interface method is the large size of the discretization matrices since the number of unknowns at the discrete level is doubled.

We demonstrated that the hexagonal channel network geometry significantly out-performs both the branching channels’ network and the classical vertical channel geometries. Our analysis indicates that the superior performance of the hexagonal geometry stems from the relatively large angle between the dominant channel flow direction and the channel-hydrogel interface. This configuration results in a larger Darcy velocity, thereby facilitating enhanced advection-mediated oxygen supply to the transplanted cells. This study is significant because recent developments in hydrogel fabrication make it now possible to control hydrogel rheology [20, 14], utilizing the computational results to generate optimized scaffold architectures.

Our future work includes the design of a geometric optimization algorithm for optimal scaffold architecture design.

## Acknowledgments

We would like to acknowledge the support by Dr. Shuvo Roy and his Bioartificial Pancreas Design Lab at UC San Francisco.

## Notes

**Funding:** This work has been supported in part by the following grants: NSF DMS-2208219 and NSF DMS-2205695 (Buka č), NSF DMS-2247000, DMS-2011319, and DMS-1853340 (Čanić), by Croatia-USA bilateral grant “The mathematical framework for the diffuse interface method applied to coupled problems in fluid dynamics” (Muha) and by the Croatian Science Foundation, project number IP-2022-10-2962 (Muha).

### Competing Interest Statement

The authors have declared no competing interest.

## REFERENCES

[1] D. M. Anderson, G. B. McFadden, and A. A. Wheeler, Diffuse-interface methods in fluid mechanics, Annual review of fluid mechanics, 30 (1998), pp. 139–165.

[2] D. N. Arnold, F. Brezzi, and M. Fortin, A stable finite element for the stokes equations, Calcolo, 21 (1984), pp. 337–344.

[3] E. S. Avgoustiniatos, K. E. Dionne, D. F. Wilson, M. L. Yarmush, and C. K. Colton, Measurements of the effective diffusion coefficient of oxygen in pancreatic islets, Industrial & engineering chemistry research, 46 (2007), pp. 6157–6163.

[4] D. Beard and J. Bassingthwaighte, Modeling advection and diffusion of oxygen in complex vascular networks, Annas of Biomedical Engineering, 29 (2001), pp. 298–310.

[5] P. Buchwald, Fem-based oxygen consumption and cell viability models for avascular pancreatic islets, Theoretical Biology and Medical Modelling, 6 (2009), pp. doi:10.1186/1742–4682–6–5.

[6] P. Buchwald, A local glucose-and oxygen concentration-based insulin secretion model for pancreatic islets, Theoretical Biology and Medical Modelling, 8 (2011), p. http://www.tbiomed.com/content/8/1/20.

[7] M. Bukač, B. Muha, and A. J. Salgado, Analysis of a diffuse interface method for the stokes-darcy coupled problem, ESAIM: Mathematical Modelling and Numerical Analysis, 57 (2023), pp. 2623–2658.

[8] M. Bukač, I. Yotov, R. Zakerzadeh, and P. Zunino, Partitioning strategies for the interaction of a fluid with a poroelastic material based on a Nitsche’s coupling approach, Computer Methods in Applied Mechanics and Engineering, 292 (2015), pp. 138–170.

[9] A.-K. Classen, K. I. Anderson, E. Marois, and S. Eaton, Hexagonal packing of Drosophila wing epithelial cells by the planar cell polarity pathway, Dev. Cell, 9 (2005), pp. 805–817.

[10] A.-K. Classen, K. I. Anderson, E. Marois, and S. Eaton, Hexagonal packing of Drosophila wing epithelial cells by the planar cell polarity pathway, Dev Cell., 9 (2005), pp. 805–17.

[11] J. Collins, A. Rudenski, J. Gibson, L. Howard, and R. O’Driscoll, Relating oxygen partial pressure, saturation and content: the haemoglobin-oxygen dissociation curve, Breathe (Sheffield, England), 11 (2015), pp. 194–201.

[12] T. Desai and L. Shea, Advances in islet encapsulation technologies, Nature Reviews (Drug Discovery), 16 (2017), pp. 338–351.

[13] H. Emmerich, The diffuse interface approach in materials science: thermodynamic concepts and applications of phase-field models, vol. 73, Springer Science & Business Media, 2003.

[14] R. S. et al., Superporous agarose scaffolds for encapsulation of adult human islets and human stem-cell-derived cells for intravascular bioartificial pancreas applications, J Biomed Mater Res, 109 (2021), pp. 2438–2448.

[15] F. H. Fenton, E. M. Cherry, A. Karma, and W.-J. Rappel, Modeling wave propagation in realistic heart geometries using the phase-field method, Chaos: An Interdisciplinary Journal of Nonlinear Science, 15 (2005), p. 013502.

[16] W. H. Fissell, A. Dubnisheva, A. N. Eldridge, A. Fleischman, A. L. Zydney, and S. Roy, High-performance silicon nanopore hemofiltration membranes, Journal of Membrane Science, 326 (2009), pp. 58–63.

[17] D. Goldman, Theoretical models of microvascular oxygen transport to tissue, Microcurculation, 15 (2008), pp. 795–811.

[18] F. Hecht, New development in FreeFem++, Journal of Numerical Mathematics, 20 (2012), pp. 251–266.

[19] D. Kanani, W. H. Fissell, S. Roy, A. Dubnisheva, A. Fleischman, and A. L. Zydney, Permeability-selectivity analysis for ultrafiltration: Effect of pore geometry, J. Memb. Sci., 349 (2010), pp. 405–418.

[20] M. Krishani, W. Y. Shin, H. Suhaimi, and N. S. Sambudi, Development of scaffolds from bio-based natural materials for tissue regeneration applications: A review, Gels, 9 (2023), p. 100.

[21] W. Martin, E. Cohen, R. Fish, and P. Shirley, Practical ray tracing of trimmed nurbs surfaces, Journal of Graphics Tools, 5 (2000), pp. 27–52.

[22] B. McGuire and T. Secomb, A theoretical model for oxygen transport in skeletal muscle under conditions of high oxygen demand, J. Appl. Physiol., 91 (2001), pp. 2255–2265.

[23] B. McGuire and T. Secomb, Estimation of capillary density in human skeletal muscle based on maximal oxygen consumption rates, Am J Physiol Heart Circ Physiol, 285 (2003), pp. H2382–H2391.

[24] O. Oyekole and M. Bukač, Second-order, loosely coupled methods for fluid-poroelastic material interaction, Numerical Methods for Partial Differential Equations, 36 (2020), pp. 800–822.

[25] A. G. Santandreu, P. Taheri-Tehrani, B. Feinberg, A. Torres, C. Blaha, R. Shaheen, J. Moyer, N. Wright, G. L. Szot, W. H. Fissell, et al., Characterization of human islet function in a convection-driven intravascular bioartificial pancreas, Bioengineering & Translational Medicine, 8 (2023), p. e10444.

[26] S. Song, C. Blaha, W. Moses, J. Park, N. Wright, J. Groszek, W. Fissell, S. Vartanian, A. M. Posselt, and S. Roy, An intravascular bioartificial pancreas device (ibap) with silicon nanopore membranes (snm) for islet encapsulation under convective mass transport, Lab. Chip, 17 (2017), pp. 1778–1792.

[27] S. Song, G. Gaetano Faleo, R. Yeung, R. Kant, A. M. Posselt, T. A. Desai, Q. Tang, and S. Roy, Silicon nanopore membrane (SNM) for islet encapsulation and immunoisolation under convective transport, Nature Scientific Reports, 6 (2016), pp. 1–9.

[28] S. K. Stoter, P. Müller, L. Cicalese, M. Tuveri, D. Schillinger, and T. J. Hughes, A diffuse interface method for the Navier–Stokes/Darcy equations: Perfusion profile for a patient-specific human liver based on MRI scans, Computer Methods in Applied Mechanics and Engineering, 321 (2017), pp. 70–102.

[29] X. Wang, Bioartificial organ manufacturing technologies, Cell Transplant, 28 (2019), pp. 5–17.

[30] Y. Wang, S. Canic, M. Bukac, C. Blaha, and S. Roy, Mathematical and computational modeling of a poroelastic cell scaffold in a bioartificial pancreas, Fluids, 7 (2022), p. 222.

[31] Y. Wang, S. Čanić, M. Bukač, C. Blaha, and S. Roy, Mathematical and computational modeling of poroelastic cell scaffolds used in the design of an implantable bioartificial pancreas, Fluids, 7 (2022), p. 222.

